# Sorcin couples Annexin A11 recruitment to ESCRT-III assembly for plasma membrane repair

**DOI:** 10.64898/2026.04.10.717788

**Authors:** Jordan Matthew Ngo, Justin Krish Williams, Abinayaa Murugupandiyan, Randy Schekman

## Abstract

The absence of a cell wall affords animal cells diverse functionality at the cost of acute sensitization to plasma membrane (PM) damage. Thus, animal cells tightly monitor and maintain PM integrity to prevent cell death. Genetic loss of PM repair factors is associated with human diseases such as muscular dystrophy. Despite evidence that annexin and endosomal sorting complex required for transport (ESCRT) proteins are required for PM repair, the extent to which their recruitment is coordinated at sites of membrane damage remains unclear. Here, leveraging quantitative organellar proteomics and genome-wide CRISPR interference screens, we identify sorcin as a new PM repair factor that couples annexin A11 (ANXA11)-mediated sensing of PM damage to ESCRT-III assembly. We show that sorcin directly binds ANXA11 and ALIX in the presence of Ca^2+^ via its penta-EF-hand domain and flexible N-terminus, respectively, and is required for ESCRT-III recruitment to PM lesions and membrane resealing. Our data support a model in which ANXA11, recruited to the PM upon damage-induced Ca^2+^ influx, serves as an anchor that facilitates the sequential recruitment of sorcin and ESCRT-III at PM lesions. Together, these findings establish a Ca^2+^-dependent scaffolding mechanism that couples PM damage sensing to ESCRT-III assembly for PM repair.

## Introduction

A defining feature of animal cells is the lack of a cell wall. Although this absence confers animal cells with distinct functional advantages compared to cells from other taxa, it also renders animal cells susceptible to plasma membrane (PM) damage. PM disruption is common, and a subset of cells, particularly those found in mechanically active tissues, are especially vulnerable to damage (1). For example, muscle and endothelial cells experience high levels of PM damage due to repeated contractions and fluid shear stress, respectively (2–4). If left unresolved, persistent PM damage leads to cell death. Accordingly, animal cells tightly monitor and maintain PM integrity.

Extracellular calcium (Ca^2+^) is essential for PM repair (5). Upon PM damage, Ca^2+^ flows from the extracellular space into the cytoplasm and initiates the recruitment of membrane repair factors to the lesion site (1). Annexins are cytosolic proteins that bind phospholipids in the presence of Ca^2+^ and participate in various aspects of PM repair, including membrane patching and tension reduction (1, 6, 7). Consistent with an important role in PM repair, genetic depletion of annexins in cultured cells impairs membrane resealing (8–12), and loss of annexin expression or function in mice causes muscular dystrophy (13–15).

In addition to annexins, the endosomal sorting complex required for transport (ESCRT) machinery promotes PM repair (16). The core ESCRT machinery, consisting of ESCRT-0, ESCRT-I, ESCRT-II, ESCRT-III, ALIX and Vps4, were originally identified for their role in sorting ubiquitylated membrane proteins into the intraluminal vesicles of multivesicular bodies (17, 18). However, it is now clear that the ESCRT machinery mediates reverse-topology membrane scission in diverse cellular processes including nuclear envelope sealing, autophagosome closure, cytokinesis, viral budding and lysosome repair (19–21). Interestingly, ESCRT I-III are recruited to damaged lysosomes (22, 23), whereas only ESCRT-III is recruited to PM lesions (24). Upon PM disruption, annexin A7 (ANXA7) localizes to the membrane lesion and recruits ALG-2, which subsequently recruits ESCRT-III (25). However, recent studies have suggested that ALG-2 can directly bind membranes in the presence of Ca^2+^ (26, 27). Thus, the extent to which annexin recruitment is coupled to ESCRT-III assembly during PM repair remains unclear.

Mutations in annexin A11 (ANXA11) and CHMP2B (a member of ESCRT-III) have been linked to amyotrophic lateral sclerosis (ALS), frontotemporal dementia (FTD) and limb-girdle muscular dystrophy (28–31), and a recent report has indicated that these mutations compromise PM repair (32). Given increasing evidence that PM repair is compromised in neurodegeneration and certain types of muscular dystrophy, we sought to employ complementary biochemical and genetic approaches to identify novel factors required for this essential membrane repair pathway.

## Results

### Identification of proteins recruited to the inner leaflet of the PM upon Ca^2+^ influx

Ca^2+^ influx upon PM disruption recruits a repertoire of membrane repair proteins to the lesion site and is required for repair (2). We sought to leverage these features of PM repair to identify new repair factors. We designed a membrane fractionation strategy that would allow us to identify proteins that are recruited to the surface of inside-out PM vesicles in the presence of Ca^2+^, reasoning that some of these factors might promote PM repair. We generated an HCT116 cell line that inducibly expressed monomeric enhanced green fluorescent protein (mEGFP) fused to the PM targeting sequence of Lck and a 3xHA epitope tag (Lck-mEGFP-3xHA) (Fig. 1*A*). Incubation of these cells with doxycycline induced titratable expression of Lck-mEGFP-3xHA (Fig. 1*B*), and visualization of these cells after doxycycline induction revealed specific labeling of the PM with mEGFP (Fig. 1*C*).

**Figure 1.**
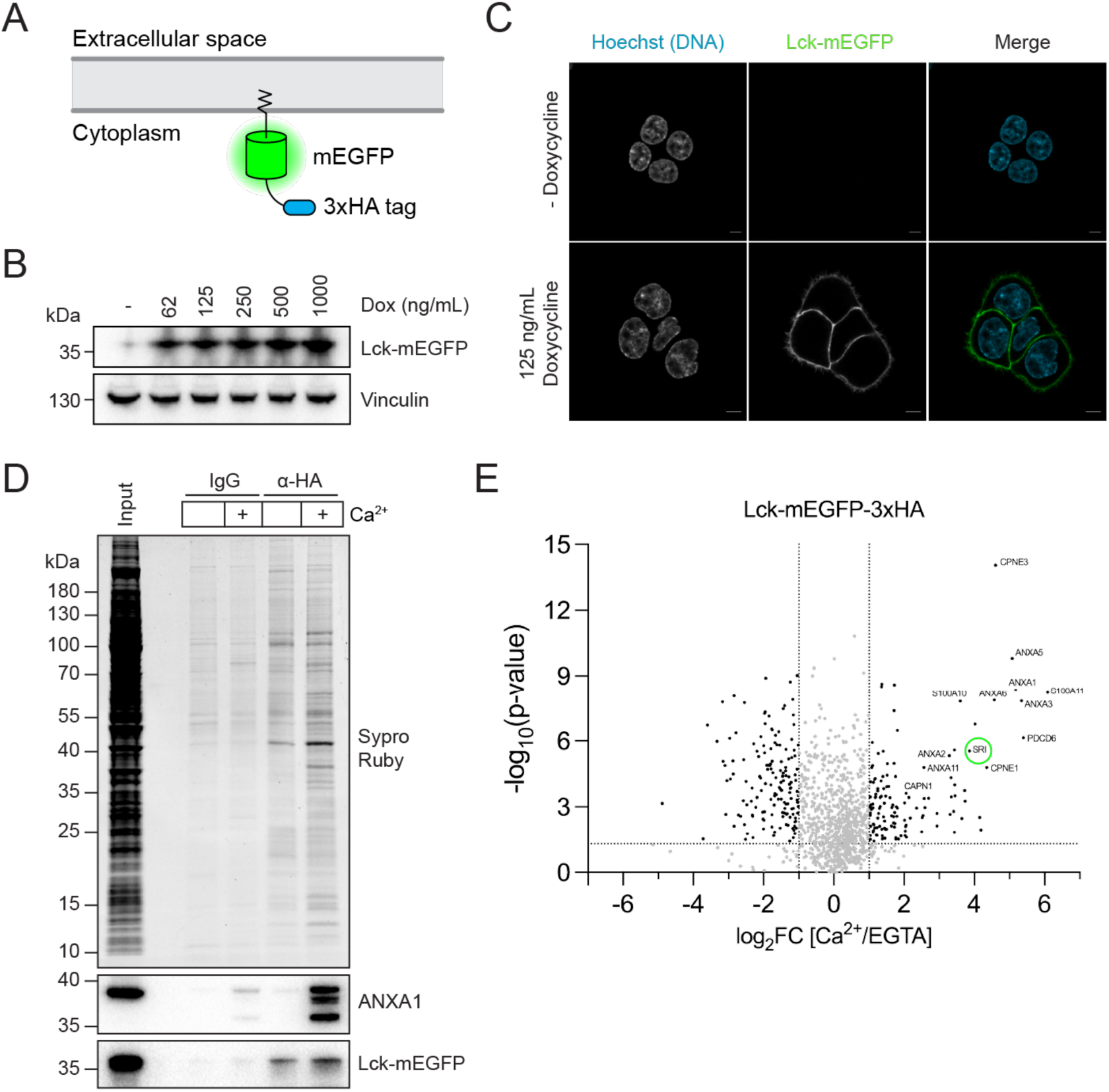
Quantitative organellar proteomics identifies Ca^2+^-dependent PM-binding proteins. (A) Schematic illustrating the membrane topology of Lck-mEGFP-3xHA when inserted into the plasma membrane. (B) Immunoblot analysis of HCT116 Lck-mEGFP-3xHA cells treated with increasing concentrations of doxycycline for 24 h. (C) Airyscan microscopy of HCT116 Lck-mEGFP-3xHA cells treated with vehicle or 125 ng/ml doxycycline for 24 h. Cyan: DNA (Hoechst 33342); Green: Lck-mEGFP-3xHA. Scale bar: 5 µm. (D) Total protein (Sypro Ruby staining) analysis of post-nuclear supernatant input and IgG and anti-HA immunoprecipitants in the absence or presence of 1 mM free Ca^2+^. (E) Volcano plot showing proteins enriched in the anti-HA immunoprecipitants in the presence of Ca^2+^. Sorcin (SRI) is circled in green. Statistical significance was calculated using an unpaired two-tailed Student’s *t*-test. See corresponding Table 1.

To identify proteins that were recruited to the PM inner leaflet upon Ca^2+^ influx, we induced Lck-mEGFP-3xHA expression, mechanically ruptured cells and performed IgG or anti-HA immunoprecipitations in the absence or presence of Ca^2+^. We observed the recruitment of multiple proteins to the surface of inside-out PM vesicles upon addition of Ca^2+^ (Fig. 1*D*). Annexin A1 (ANXA1) served as a positive control and was greatly enriched in the anti-HA + Ca^2+^ immunoprecipitant. Label-free quantitative mass spectrometry analysis of the anti-HA immunoprecipitants identified many proteins that were reproducibly recruited to the inner leaflet of the PM in the presence of Ca^2+^ (Fig. 1*E*; Table 1). Many significant hits identified in our mass spectrometry analysis (such as annexins, S100A10/11, ALG-2 and calpains) have been previously implicated in PM repair. These data confirmed that our inside-out PM vesicle isolation strategy faithfully recapitulated the Ca^2+^-dependent recruitment of repair factors to the PM inner leaflet, validating its utility for identifying new PM repair factors.

### Genome-wide CRISPR interference screens for PM repair

Simultaneously, we performed complementary genetic screens to identify genes whose depletion sensitized cells to PM damage inflicted by treatment with the pore-forming toxin perfringolysin O (PFO). To achieve this, we first purified recombinant PFO from bacterial cells (Fig. 2*A*). Consistent with a requirement for extracellular Ca^2+^ in PM repair, an approximately 5-fold lower PFO dose was sufficient to kill cells in the absence of extracellular Ca^2+^ (Fig. 2*B*). Exploiting this sensitization, we designed a genome-wide CRISPR interference (CRISPRi) survival screen strategy to distinguish likely PM repair factors from genes that affect PFO binding (Fig. 2*C*). Library-transduced cells were treated with PFO at doses that killed 25% of cells in the presence or absence of extracellular Ca^2+^ (Fig. 2*B*). CRISPRi guide RNA (sgRNA) enrichment was determined by deep sequencing of PCR-amplified barcodes (Table 2) (33).

**Figure 2.**
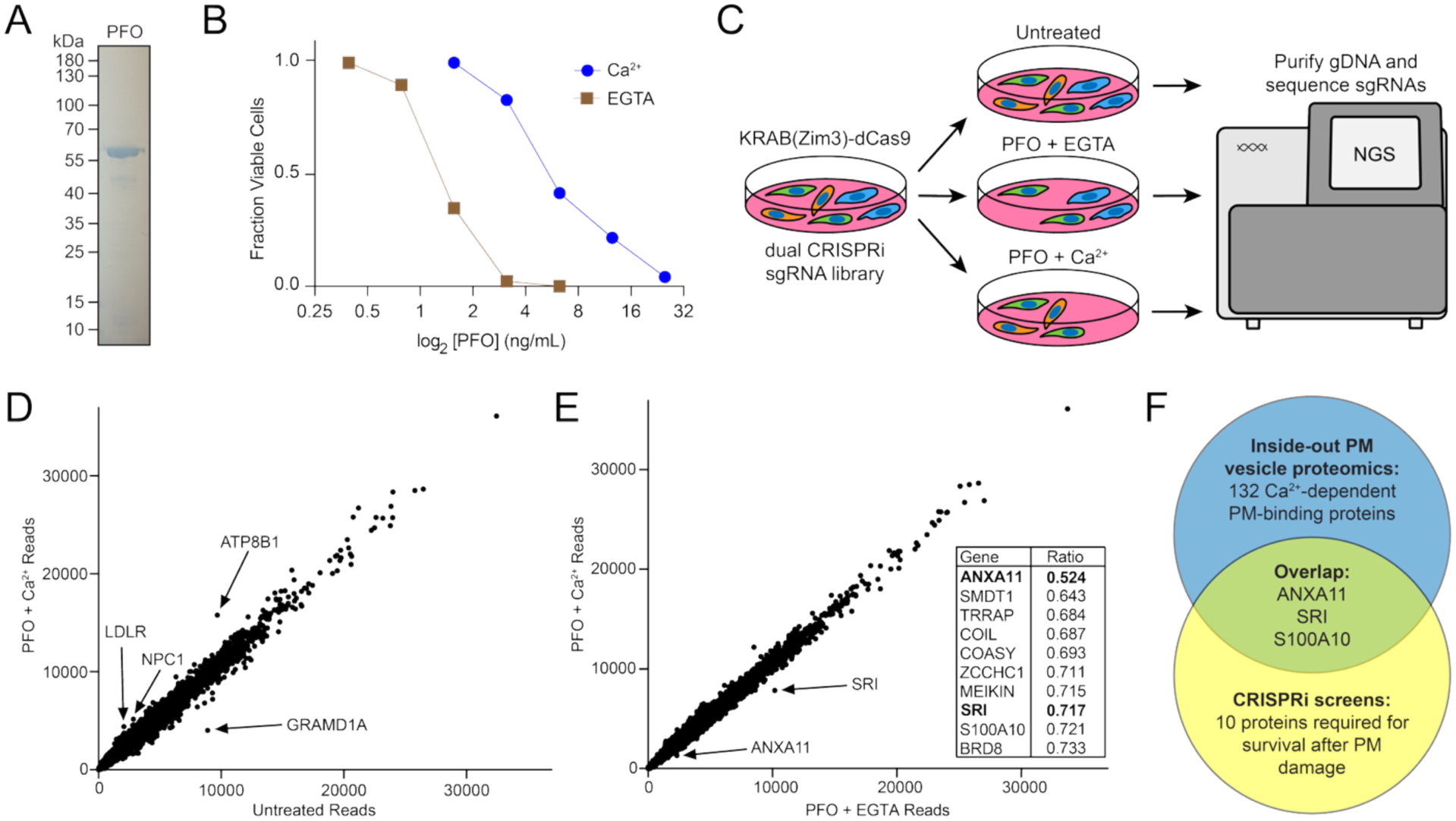
Genome-wide CRISPR interference screens identify genes required for survival after PM damage. (A) Coomassie-stained SDS-PAGE gel showing the purification of recombinant perfringolysin O (PFO). (B) Quantification of cell viability, as measured by trypan blue exclusion, plotted as a function of PFO concentration. (C) Schematic of the three genome-wide CRISPR interference (CRISPRi) screen conditions. PFO doses that killed 25% of cells were used for the genetic screens. (D) Gene enrichment comparing untreated (2 mM extracellular Ca^2+^) and PFO + 2 mM extracellular Ca^2+^ screens revealed factors involved in PFO binding and cholesterol trafficking. See corresponding Table 2. (E) Gene enrichment comparing PFO + 2 mM extracellular EGTA and PFO + 2 mM extracellular Ca^2+^ screens identified Ca^2+^-dependent plasma membrane repair (PM) factors. Inset: top 10 genes depleted under PFO + 2 mM extracellular Ca^2+^ treatment. See corresponding Table 2. (F) Venn diagram showing common hits between the inside-out PM vesicle proteomics (Figure 1E) and the CRISPRi screens (Figure 2E).

When comparing the untreated and PFO + Ca^2+^ screens, the most enriched CRISPRi sgRNAs corresponded to genes involved in PFO binding and cholesterol trafficking (Fig. 2*D*). PFO is a cholesterol-dependent cytolysin that recognizes cholesterol in the outer leaflet of the PM (34). In contrast, comparison between the PFO + EGTA (Ca^2+^ chelator) and the PFO + Ca^2+^ screens revealed several Ca^2+^-binding proteins (Fig. 2*E*, Inset).

Comparison of the hits identified in our proteomic and genetic screens revealed 3 overlapping genes: annexin A11 (ANXA11), sorcin (SRI), and S100A10 (Fig. 2*F*). ANXA11 and S100A10 have been previously implicated in PM repair. ALS-linked mutations in ANXA11 affect ESCRT-III dynamics at the PM (32). We previously showed that a S100A10-annexin A2 (ANXA2) complex localizes to PM lesions before being cleaved by calpains (35). Of particular interest, sorcin emerged as a completely novel hit identified in both our proteomic and genetic screens (Fig. 2*F*).

### Sorcin is recruited to PM lesions upon Ca^2+^ influx and required for membrane repair

Sorcin, like ALG-2, is a member of the Ca^2+^-binding penta-EF-hand (PEF) family of proteins (36). Sorcin was initially identified in multidrug-resistant cancer cells and has been suggested to regulate intracellular Ca^2+^ homeostasis (37–42). Despite this, a role for sorcin in membrane repair has not been described. To determine whether sorcin was a bona fide PM repair factor, we first tested whether sorcin is recruited to PM lesions. We generated a U2-OS cell line expressing fluorescently tagged sorcin (SRI-mNG) to monitor sorcin dynamics after laser ablation wounding. FM4-64 is a membrane-impermeable dye that brightly labels the site of damage and the PM repair cap. Upon laser ablation, we observed FM4-64 staining and sorcin recruitment at the site of PM damage (Fig. 3*A*; Movie 1). We next tested whether endogenous sorcin is recruited to PM lesions. Immunofluorescence microscopy of vehicle-and PFO-treated cells revealed that endogenous sorcin is recruited to sites of PM damage (Fig. 3*B*). Because extracellular Ca^2+^ is essential for PM repair (5), we asked whether Ca^2+^ influx was required for sorcin recruitment to PM lesions. Laser ablation experiments revealed that the removal of extracellular Ca^2+^ completely blocked both PM repair cap formation and sorcin recruitment to PM lesions (Fig. 3*C*; Movies 2 and 3).

**Figure 3.**
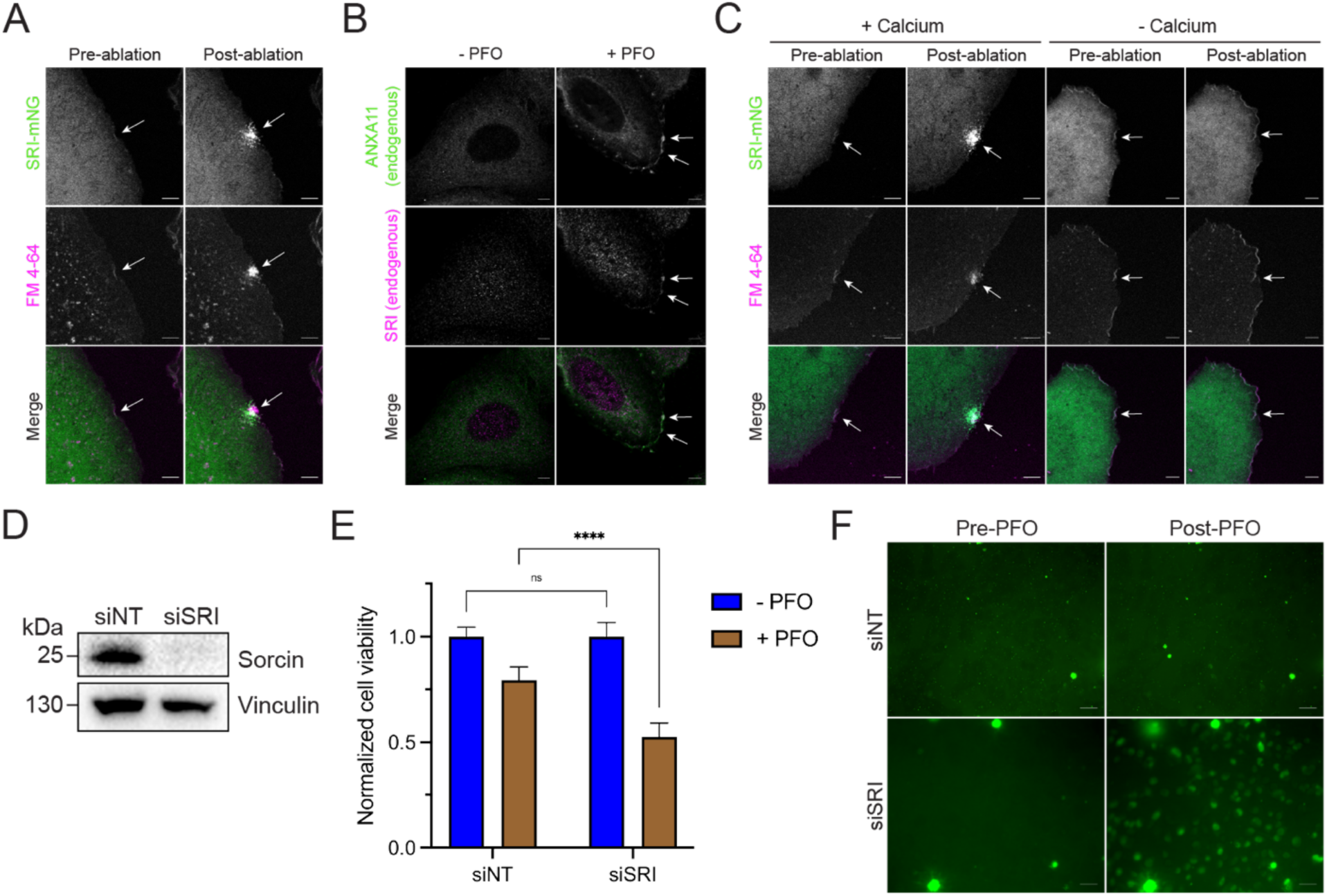
Sorcin is recruited to PM lesions upon Ca^2+^ influx and required for membrane repair. (A) Representative confocal micrographs of U2-OS Sorcin-mNeonGreen (SRI-mNG) cells subjected to laser ablation wounding. Images shown are pre-and post-laser ablation. Green: SRI-mNG; Magenta: FM4-64. Scale bar: 5 µm. See corresponding Movie 1. (B) Immunofluorescence microscopy of endogenous ANXA11 and SRI from U2-OS cells treated with vehicle or perfringolysin O (PFO; 25 ng/ml) for 5 min. Green: ANXA11; Magenta: SRI. Scale bar: 5 µm. (C) Representative confocal micrographs of U2-OS SRI-mNG cells subjected to laser ablation wounding in the presence or absence of 2 mM extracellular Ca^2+^. Images shown are pre-and post-laser ablation. Green: SRI-mNG; Magenta: FM4-64. Scale bar: 5 µm. See corresponding Movies 2 and 3. (D) Immunoblot analysis of U2-OS wild-type cells transfected with control (siNT) or sorcin-targeting (siSRI) siRNAs. (E) Normalized cell viability of U2-OS siNT and siSRI cells following treatment with vehicle or PFO (25 ng/ml). Graph displays mean ± SD (n = 9). Statistical significance was calculated using a two-way ANOVA (ns = not significant, ****p<0.0001). (F) Representative confocal micrographs of U2-OS siNT and siSRI cells treated with PFO (25 ng/ml) in the presence of SYTOX Green (2.5 µM). Images shown are pre-and post-PFO treatment. Green: SYTOX Green. Scale bar: 50 µm.

We then asked whether sorcin depletion sensitized cells to PM damage. To address this, we depleted sorcin by RNA interference (Fig. 3*D*) and treated control (siNT) and sorcin-depleted (siSRI) cells with PFO (Fig. 3*E*). We observed that sorcin depletion sensitized cells to PFO treatment. We then tested whether sorcin depletion inhibited PM repair by monitoring the influx of the membrane-impermeable dye SYTOX Green into siNT and siSRI cells upon PFO treatment. Measurement of SYTOX Green influx revealed that sorcin depletion compromised PM repair (Fig. 3*F*). Together, these experiments established sorcin as a bona fide PM repair factor.

### Sorcin simultaneously binds ANXA11 and ALIX in the presence of Ca^2+^

Prompted by our observation that Ca^2+^ influx is essential for sorcin recruitment to PM lesions (Fig. 3*C*), we sought to identify proteins that interact with sorcin in the presence of Ca^2+^. Sorcin was previously reported to interact with a recombinant N-terminal fragment of ANXA11 (43), the strongest Ca^2+^-dependent candidate identified in our CRISPRi screens for cell survival upon PFO challenge (Fig. 2*E*). Additionally, ALG-2 has been demonstrated to bind ALIX (44). Motivated by these observations, we hypothesized that sorcin associates with ANXA11 and components of the ESCRT-III complex in the presence of Ca^2+^.

To test this hypothesis, we first performed immunoprecipitation experiments to evaluate whether endogenous sorcin associates with ANXA11 and ALIX. Immunoblot analysis of IgG and anti-sorcin immunoprecipitants revealed that endogenous sorcin associates with ANXA11 and ALIX in the presence of Ca^2+^ (Fig. 4*A*). We then sought to determine which domain of sorcin mediates its interaction with ANXA11 and ALIX. Alphafold3 modeling revealed that sorcin consists of two domains: a flexible N-terminal domain (amino acids 1-32) and a PEF domain (amino acids 33-198) (Fig. 4*B*). Given previous evidence indicating that the N-terminus of PEF proteins regulates their association with other proteins (45), we purified FLAG-tagged full-length (SRI^FL^-FLAG) and truncated (SRI^Δ1-32^-FLAG) sorcin from bacterial cells (Fig. 4 *C* and *D*) for immunoprecipitation experiments. Full-length and truncated sorcin were immobilized on anti-FLAG beads, incubated with cytosol in the absence or presence of Ca^2+^, and analyzed by immunoblotting (Fig. 4*E*). These experiments revealed that sorcin bound to ANXA11 and ALIX in the presence of Ca^2+^. Notably, truncated sorcin retained its ability to bind ANXA11, but not ALIX. We then asked whether sorcin directly associates with ALIX. I*n vitro* binding assays with purified recombinant proteins revealed that sorcin directly binds ALIX in the presence of Ca^2+^ (Fig. 4*F*). Together, these data demonstrated that sorcin simultaneously binds ANXA11 and ALIX in the presence of Ca^2+^.

**Figure 4.**
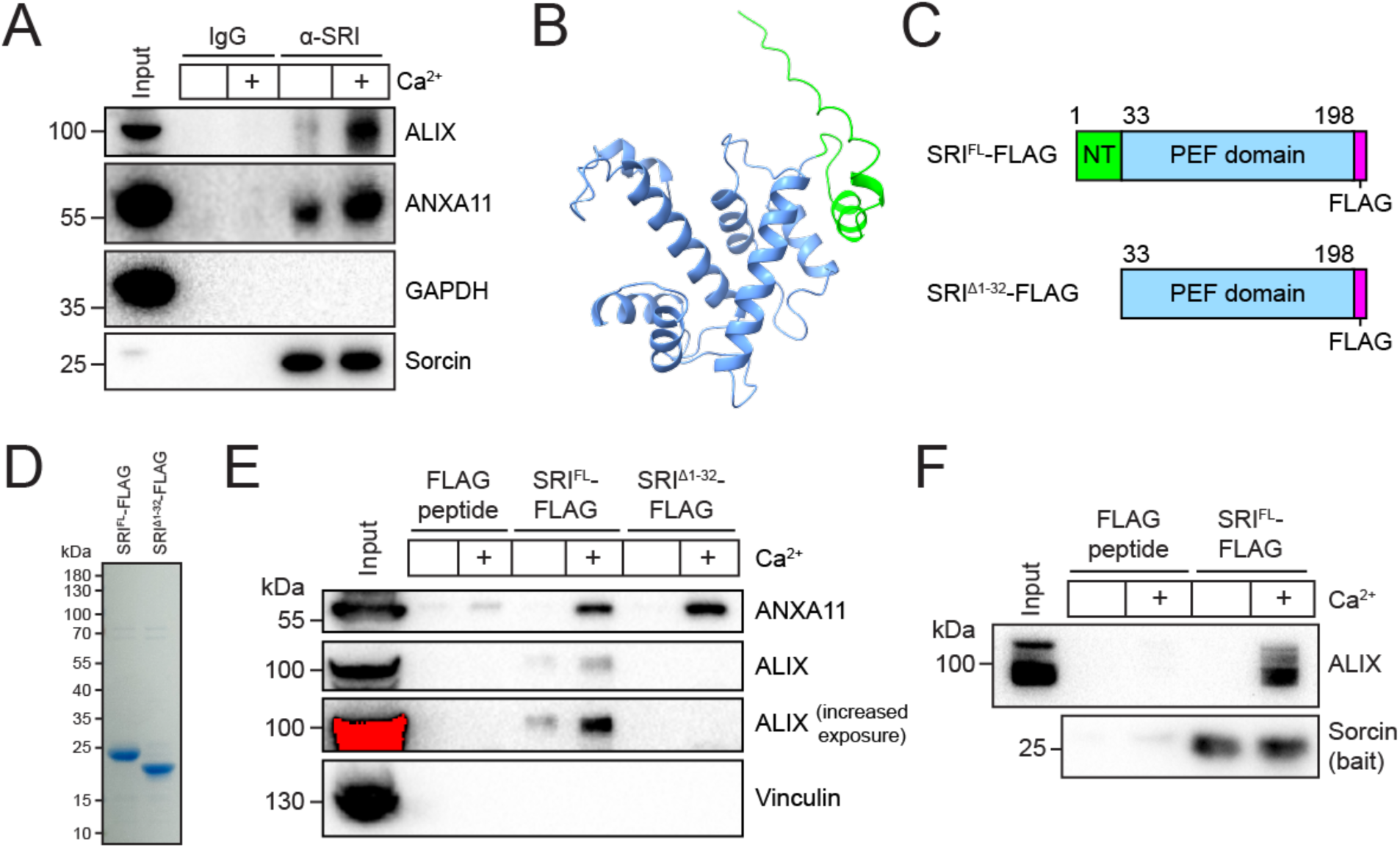
**Sorcin simultaneously binds ANXA11 and ALIX in the presence of Ca^2+^.**(A) Immunoblot analysis of post-nuclear supernatant input and IgG and anti-sorcin immunoprecipitants in the absence or presence of 1 mM free Ca^2+^. (B) Alphafold3 predicted protein structure of human sorcin (SRI). The flexible N-terminal domain (NT) is colored green, and the penta-EF hand (PEF) domain is colored blue. (C) Domain architecture of full-length (SRI^FL^-FLAG) and truncated (SRI^Δ1-32^-FLAG) sorcin constructs. (D) Coomassie-stained SDS-PAGE gel showing the purification of SRI^FL^-FLAG and SRI^Δ1-32^-FLAG. (E) Immunoblot analysis of cytosol input and FLAG peptide, SRI^FL^-FLAG and SRI^Δ1-32^-FLAG immunoprecipitants in the absence or presence of 1 mM free Ca^2+^. (F) Immunoblot analysis of recombinant protein input and immunoprecipitants from an SRI^FL^-FLAG/ALIX *in vitro* binding assay in the absence or presence of 1 mM free Ca^2+^.

### ANXA11 is required for membrane repair and sorcin recruitment to PM lesions

Our sorcin truncation data led us to hypothesize that sorcin couples ANXA11 recruitment and ESCRT-III assembly during PM repair. We thus evaluated the role of ANXA11 in PM repair and sorcin recruitment to PM lesions. We first tested whether ANXA11 was required for PM repair. We depleted ANXA11 by RNA interference (Fig. 5*A*) and treated siNT and ANXA11-depleted (siANXA11) cells with PFO (Fig. 5*B*). This experiment revealed that ANXA11 depletion sensitized cells to PFO. We then performed SYTOX Green influx experiments in siNT and siANXA11 cells and found that ANXA11 depletion compromised PM repair (Fig. 5*C*).

**Figure 5.**
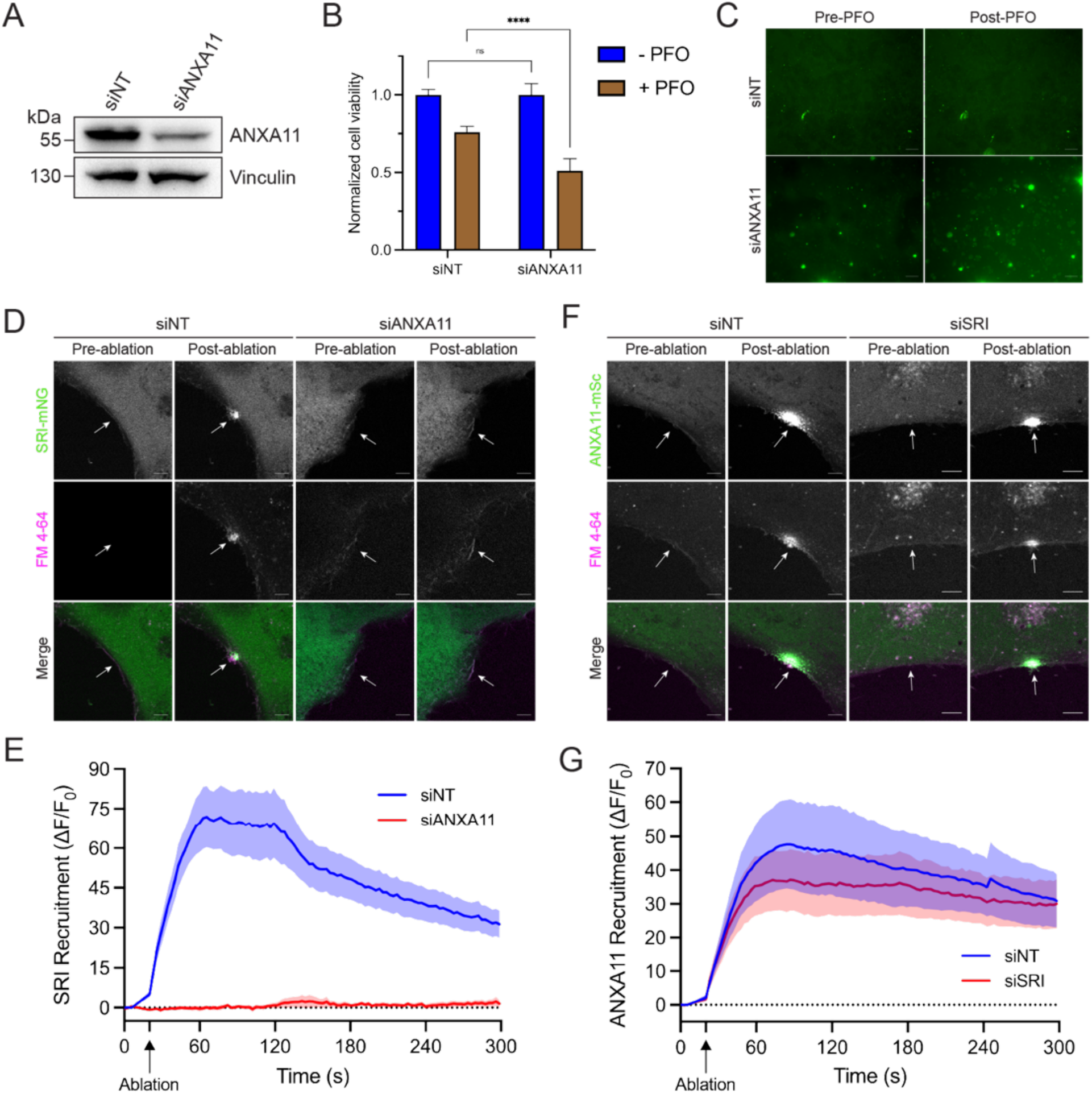
ANXA11 is required for PM repair and sorcin recruitment to PM lesions. (A) Immunoblot analysis of U2-OS wild-type cells transfected with control (siNT) or ANXA11-targeting (siANXA11) siRNAs. (B) Normalized cell viability of U2-OS siNT and siANXA11 cells following treatment with vehicle or perfringolysin O (PFO; 25 ng/ml). Graph displays mean ± SD (n = 6). Statistical significance was calculated using a two-way ANOVA (ns = not significant, ****p<0.0001). (C) Representative confocal micrographs of U2-OS siNT and siANXA11 cells treated with PFO (25 ng/ml) in the presence of SYTOX Green (2.5 µM). Images shown are pre-and post-PFO treatment. Green: SYTOX Green. Scale bar: 50 µm. (D) Representative confocal micrographs of U2-OS Sorcin-mNeonGreen (SRI-mNG) siNT and siANXA11 cells subjected to laser ablation wounding. Images shown are pre-and post-laser ablation. Green: SRI-mNG; Magenta: FM4-64. Scale bar: 5 µm. See corresponding Movies 4 and 5. (E) Quantification of the recruitment dynamics of SRI-mNG cells transfected with siNT (n = 19) or siANXA11 (n = 14) siRNAs. Graph displays mean ± S.E.M. (F) Representative confocal micrographs of U2-OS ANXA11-mScarlet (ANXA11-mSc) siNT and sorcin knockdown (siSRI) cells subjected to laser ablation wounding. Images shown are pre-and post-laser ablation. Green: ANXA11-mSc; Magenta: FM4-64. Scale bar: 5 µm. See corresponding Movies 6 and 7. (G) Quantification of the recruitment dynamics of ANXA11-mSc cells transfected with siNT (n = 9) or siSRI (n = 13) siRNAs. Graph displays mean ± S.E.M.

We then asked whether ANXA11 is required for sorcin recruitment to PM lesions. Laser ablation studies revealed that ANXA11 depletion blocked sorcin recruitment to PM lesions (Fig. 5 *D* and *E;* Movies 4 and 5). In contrast, sorcin depletion did not affect ANXA11 recruitment to PM lesions (Fig. 5 *F* and *G;* Movies 6 and 7). Notably, PM repair cap formation (as indicated by FM4-64 staining) was defective in ANXA11-depleted cells (Fig. 5*D*).

### Sorcin and ANXA11 are required for ESCRT-III recruitment to PM lesions

After establishing roles for ANXA11 in PM repair and sorcin recruitment to PM lesions, we asked whether sorcin and ANXA11 were required for ESCRT-III recruitment. To address this, we depleted sorcin or ANXA11 in U2-OS cells expressing fluorescent ALIX and CHMP4B fusion proteins and performed laser ablation studies. Strikingly, depletion of either sorcin or ANXA11 blocked the recruitment of both ALIX (Fig. 6 *A* and *B*; Movies 8-10) and CHMP4B (Fig. 6 *C* and *D*; Movies 11-13) to PM lesions.

**Figure 6.**
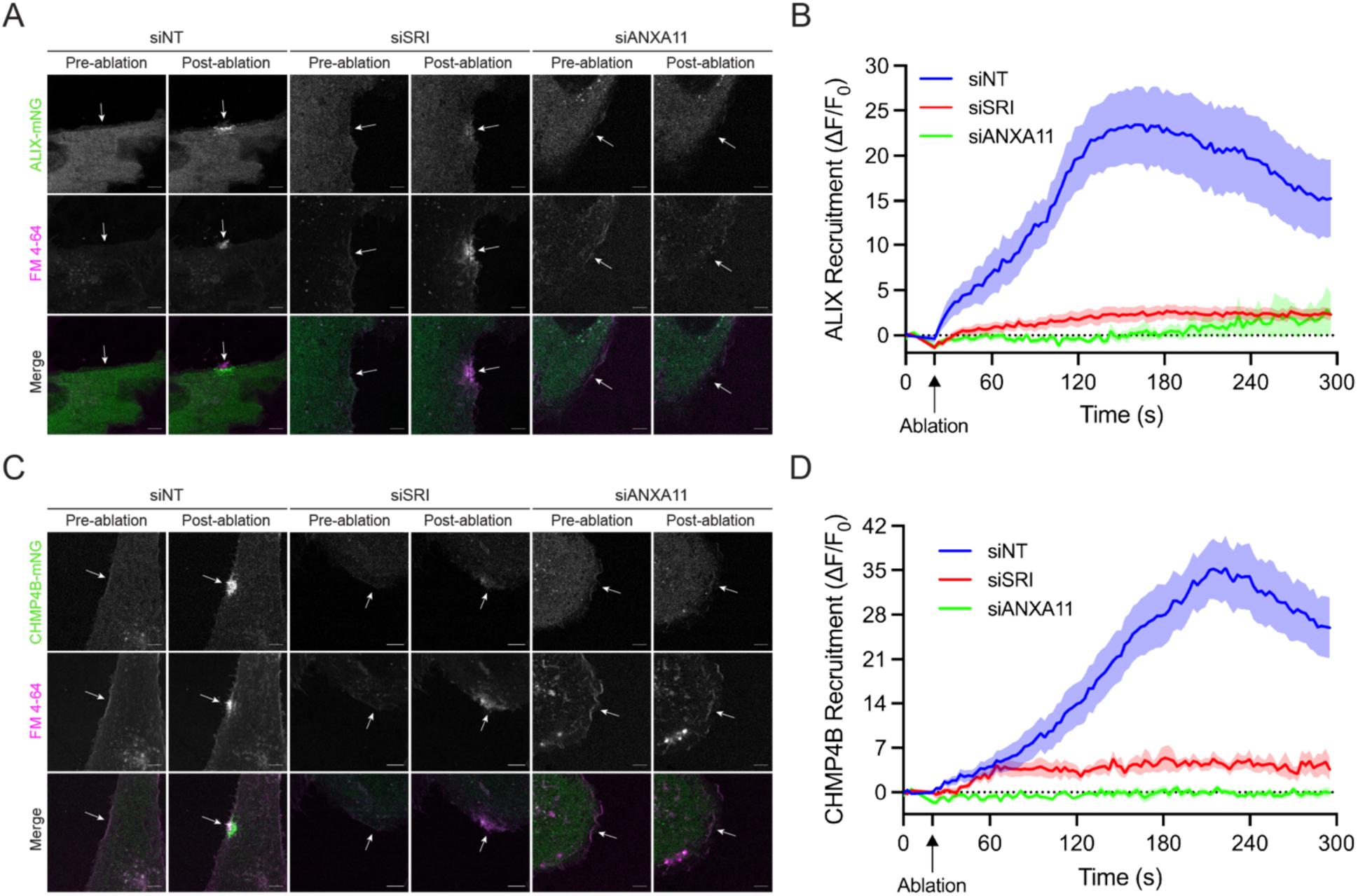
Sorcin and ANXA11 are required for ESCRT-III recruitment to PM lesions. (A) Representative confocal micrographs of U2-OS ALIX-mNeonGreen (ALIX-mNG) control (siNT), sorcin knockdown (siSRI) and ANXA11 knockdown (siANXA11) cells subjected to laser ablation wounding. Images shown are pre-and post-laser ablation. Green: ALIX-mNG; Magenta: FM4-64. Scale bar: 5 µm. See corresponding Movies 8-10. (B) Quantification of the recruitment dynamics of ALIX-mNG cells transfected with siNT (n = 15), siSRI (n = 12) or siANXA11 (n = 11) siRNAs. Graph displays mean ± S.E.M. (C) Representative confocal micrographs of U2-OS CHMP4B-mNeonGreen (CHMP4B-mNG) siNT, siSRI and siANXA11 cells subjected to laser ablation wounding. Images shown are pre-and post-laser ablation. Green: CHMP4B-mNG; Magenta: FM4-64. Scale bar: 5 µm. See corresponding Movies 11-13. (D) Quantification of the recruitment dynamics of CHMP4B-mNG cells transfected with siNT (n = 15), siSRI (n = 7) or siANXA11 (n = 7) siRNAs. Graph displays mean ± S.E.M.

### Sorcin is recruited to PM lesions together with ANXA11 and before ESCRT-III

We then sought to establish the chronology in which ANXA11, sorcin, ALIX and CHMP4B are recruited to sites of PM damage. We re-analyzed our laser ablation data (Figs. 5 and 6) and quantified the recruitment kinetics of each protein at PM lesions (Fig. 7 *A* and *B*). This analysis revealed that ANXA11 and sorcin are simultaneously recruited to PM lesions, followed sequentially by ALIX and then CHMP4B. These data, taken together with our truncation (Fig. 4) and laser ablation (Figs. 5 and 6) data, suggested a model in which ANXA11 anchors at PM lesions to facilitate sorcin recruitment and subsequent ESCRT-III assembly for PM repair.

**Figure 7.**
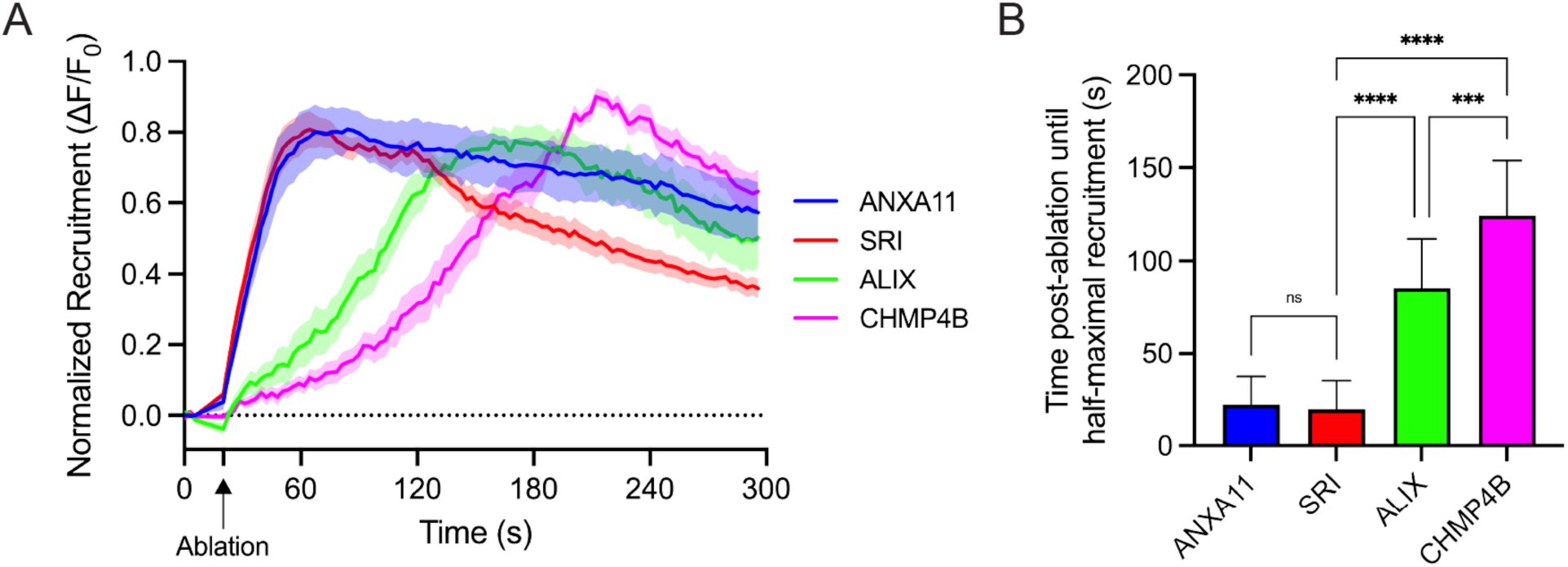
Sorcin is recruited to PM lesions together with ANXA11 and before ESCRT-III. (A) Quantification of the recruitment dynamics of ANXA11-mSc (n = 9), Sorcin-mNG (n = 19), ALIX-mNG (n = 15) and CHMP4B-mNG (n = 15) cells transfected with a non-targeting siRNA (siNT) from Figures 5E, 5G, 6B and 6D. Graph displays mean ± S.E.M. (B) Quantification of the time post-ablation until half-maximal recruitment of ANXA11-mSc (n = 9), Sorcin-mNG (n = 19), ALIX-mNG (n = 15) and CHMP4B-mNG (n = 15) siNT cells from Figures 5E, 5G, 6B and 6D. Graph displays mean ± SD. Statistical significance was calculated using a one-way ANOVA (ns = not significant, ***p<0.001, ****p<0.0001).

### ANXA11 is sufficient to recruit sorcin to liposomes in the presence of Ca^2+^

We directly tested a part of this model by performing liposome flotation assays with purified components (Fig. 8*A*). Recombinant sorcin was incubated with synthetic liposomes and Ca^2+^ in the absence or presence of recombinant ANXA2 or ANXA11. The binding reactions were applied to the bottom of sucrose density gradients followed by high-speed centrifugation. Buoyant liposomes and bound proteins were solubilized then evaluated by immunoblot analysis. Sorcin alone was unable to float with liposomes, indicating that sorcin cannot directly bind membranes (Fig. 8*B*). However, sorcin floated with liposomes when ANXA11, but not ANXA2, was added to the binding reaction. These results indicated that sorcin binds directly and specifically to ANXA11 and that ANXA11 is sufficient to recruit sorcin to membranes in the presence of Ca^2+^ (Fig. 8*B*).

**Figure 8.**
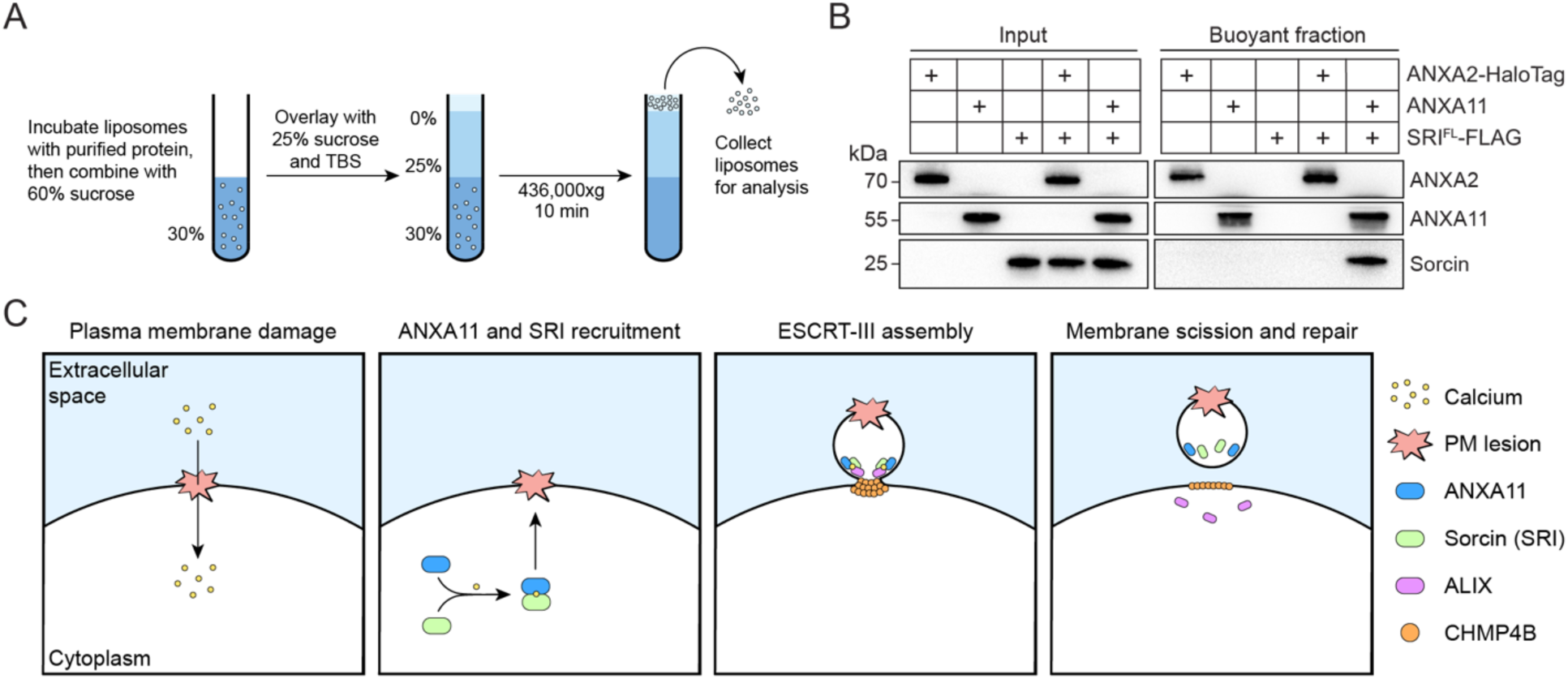
ANXA11 is sufficient to recruit sorcin to liposomes in the presence of Ca^2+^. (A) Schematic of the liposome flotation assay. Purified proteins were incubated with synthetic liposomes and 2 mM free Ca^2+^ prior to flotation through a sucrose density gradient to determine which proteins associated with membranes. (B) Immunoblot analysis of the liposome flotation assays. The components of each reaction are indicated. (C) Schematic depicting the current working model of sorcin-mediated plasma membrane repair.

## Discussion

The molecular basis by which PM damage sensing is coupled to ESCRT-III assembly remains incompletely understood. Here, we identify sorcin as an early-acting Ca^2+^-dependent scaffold that bridges ANXA11-mediated PM damage sensing and ESCRT-III assembly at sites of membrane damage (Fig. 8*C*). Our data support a model in which ANXA11, recruited to the PM upon damage-induced Ca^2+^ influx, serves as an anchor that promotes the sequential recruitment of sorcin and ESCRT-III at PM lesions (Figs. 5-8). The direct and simultaneous engagement of sorcin with ANXA11 and ALIX via its PEF domain and flexible N-terminus, respectively, provides a molecular explanation for how annexin recruitment is coupled to ESCRT-III assembly during PM repair (Fig. 4*E*). Genetic depletion of sorcin sensitizes cells to PM damage and compromises ESCRT-III recruitment, but not ANXA11 recruitment, to PM lesions (Figs. 3 *D*-*F*, 5 *F* and *G,* and 6), establishing sorcin as a critical PM repair factor.

ANXA11 depletion substantially decreases the size of the FM4-64-positive repair cap (Fig. 5*D*), consistent with our previous finding that ANXA2 depletion has a similar effect (35). Together, these data suggest that annexins play nonredundant roles in repair cap formation and that the proper balance of annexins is required for successful repair cap assembly. Furthermore, sorcin binds to the proline-rich ANXA11 N-terminus, which resembles the motif that we previously found to be cleaved by calpains during PM repair. Thus, calpains may regulate ANXA11-mediated ESCRT-III recruitment.

The ANXA11/SRI pathway may synergize with the reported ANXA7/ALG-2 pathway to recruit ESCRT-III to PM lesions (25). However, we have found some biochemical differences between sorcin and ALG-2. Our data indicate that sorcin simultaneously engages ANXA11 and ALIX via its flexible N-terminus and PEF domain, respectively. Additionally, sorcin is unable to directly bind membranes in the presence of Ca^2+^. In contrast, ALG-2 interacts with both ANXA7 and ALIX via its PEF domain (46, 47) and has recently been reported to directly bind membranes (26, 27). Thus, our data establish a mechanism by which annexin recruitment is directly coupled to ESCRT-III assembly during PM repair.

Genetic mutations in ESCRT machinery components have been implicated in various neurodegenerative diseases and membrane repair disorders (28, 48–52). In addition, ANXA11 mutations have been associated with ALS and muscular dystrophy (29–31). Although these mutations have been studied primarily in the context of lysosomal membrane repair, a recent report found that ALS-and FTD-associated mutations in ANXA11 and CHMP2B also compromise PM repair (32). Our identification of sorcin as a Ca^2+^-dependent scaffold that recruits ESCRT-III to PM lesions via ANXA11 raises the possibility that disease-associated ANXA11 mutations might impair PM repair, at least in part, by disrupting sorcin recruitment to PM lesions. Consistent with this suggestion, multiple disease-associated ANXA11 mutations map to the same flexible N-terminal region that binds sorcin.

Finally, our data highlight mechanistic and topological similarities between the budding of membrane-enveloped viruses (e.g. Human Immunodeficiency Virus [HIV-1]) and damage-induced PM vesicles (microvesicles). Göttlinger and colleagues provided the initial piece of evidence that host factors contribute to virus budding (53). Subsequent work by the Sundquist and Göttlinger teams revealed that the p6 region of the HIV-1 Gag protein engages TSG101 and ALIX, which subsequently recruit ESCRT-III and Vps4 for membrane constriction and scission (54–56). More recently, we found that PM damage elicits extracellular vesicle secretion and that expression of a dominant-negative Vps4 mutant, which blocks ESCRT-III disassembly and vesicle scission, reduces the secretion of damage-induced microvesicles (35, 57, 58). Our identification of sorcin as a membrane repair factor that localizes to PM lesions and recruits ESCRT-III for microvesicle shedding leads us to propose that enveloped viruses and PM damage-induced microvesicles share a common mechanism of membrane budding. Specifically, we propose that ANXA11 and sorcin function analogously to retroviral Gag proteins in recruiting ALIX and ESCRT-III to the PM for membrane budding. We speculate that membrane-enveloped viruses may have co-opted this host pathway of PM ESCRT recruitment to facilitate virion assembly and propagation.

## Materials and Methods

### Cell lines, media and general chemicals

HEK293T, HCT116, MDA-MB-231 and U2-OS cell lines were cultured in 5% CO_2_ at 37°C and maintained in Dulbecco’s Modified Eagle Medium (DMEM) supplemented with 10% fetal bovine serum (FBS). Cells were routinely tested, and found negative, for mycoplasma contamination using the MycoAlert Mycoplasma Detection Kit (Lonza Biosciences). All cell lines used in this study were authenticated by the UC Berkeley Cell Culture Facility using STR profiling. Unless otherwise noted, all chemicals were purchased from Sigma-Aldrich.

### Antibodies

Primary antibodies used for immunoblotting (1:1000 dilution) were: mouse anti-ALIX (Santa Cruz; sc-53540), rabbit anti-ALIX (Proteintech; 12422-1-AP), rabbit anti-annexin A1 (Abcam; ab214486), rabbit anti-annexin A2 (Abcam; ab185957), rabbit anti-annexin A11 (Proteintech; 10479-2-AP), rabbit anti-GAPDH (Cell Signaling Technology; 14C10) rabbit anti-HA-Tag (Cell Signaling Technology; C29F4), rabbit anti-SRI (Abcam; ab71983), and rabbit anti-vinculin (Abcam; ab129002). Secondary antibodies used for immunoblotting (1:10,000 dilution) were HRP-linked sheep anti-mouse IgG (Cytiva; NA931) and HRP-linked donkey anti-rabbit IgG (Cytiva; NA934).

Primary antibodies used for immunofluorescence (1:100 dilution) were mouse anti-sorcin (Santa Cruz; sc-100859) and rabbit anti-annexin A11 (Proteintech; 10479-2-AP). Secondary antibodies used for immunofluorescence (1:500 dilution) were Alexa Fluor 488 donkey anti-rabbit IgG (A-21202; Invitrogen) and Alexa Fluor 555 goat anti-mouse IgG (A-21424; Invitrogen).

### Plasmids

Lck-mEGFP-3xHA was cloned into a pLIX403-Puro backbone. mNeonGreen and mScarlet fusion proteins were cloned into pLJM1-Puro and pLJM1-Blast backbones, respectively. Constructs for recombinant protein purification were cloned into a pET28a backbone. Recombinant PFO was appended at the C-terminus with a His_6_ tag. Recombinant full-length and truncated sorcin were appended at the C-terminus with a FLAG tag and a His_6_ tag. Recombinant ANXA2 was appended at the C-terminus with HaloTag and a His_6_ tag. All plasmid constructs were verified by whole-plasmid sequencing (ElimBio).

### Lentivirus production and transduction

HEK293T cells at ∼40% confluence in a 6-well plate were transfected with 165 ng of pMD2.G, 1.35 µg of psPAX2 and 1.5 µg of a lentiviral plasmid using 7.5 µl of TransIT-LT1 Transfection Reagent (Mirus Bio) according to the manufacturer’s protocol. For lentiviral expression of the genome-wide CRISPRi library, HEK293T cells at similar confluence within two 15 cm dishes were each transfected with 1 µg of pMD2.G, 7 µg of psPAX2, and 8 µg of the Dual sgRNA CRISPRi library (Addgene #187246) using 48 µl of TransIT-LT1 Transfection Reagent. The lentivirus-containing medium was harvested 48 h post-transfection by filtration through a 0.45 μm polyethersulfone filter (VWR Sciences). The filtered lentivirus was aliquoted, snap-frozen in liquid nitrogen and stored at-80°C until use. Cells were transduced with filtered lentivirus in the presence of 8 μg/ml polybrene for 24 h after which the medium was replaced. Cells were selected using 1 μg/ml puromycin, 5 µg/ml blasticidin, or 100 µg/ml hygromycin B.

### Immunoblotting

Cells were lysed in Tris-buffered saline (TBS) supplemented with 1% Triton X-100 (TX-100) and a protease inhibitor cocktail (1 mM 4-aminobenzamidine dihydrochloride, 1 µg/ml antipain dihydrochloride, 1 µg/ml aprotinin, 1 µg/ml leupeptin, 1 µg/ml chymostatin, 1 mM phenylmethylsulfonyl fluoride (PMSF), 50 µM N-tosyl-L-phenylalanine chloromethyl ketone and 1 µg/ml pepstatin) and incubated on ice for 15 min. Whole cell lysates were centrifuged at 15,000xg for 10 min at 4°C, and the post-nuclear supernatant (PNS) was diluted with 6X Laemmli buffer (reducing) to a 1X final concentration. Samples were heated at 95°C for 5 min and resolved in 4-20% acrylamide Tris-glycine gradient gels (Life Technologies). The resolved proteins were then transferred to polyvinylidene difluoride (PVDF) membranes (EMD Millipore).

The PVDF membranes were blocked for 30 min with 5% bovine serum albumin (BSA) in TBS supplemented with 0.1% Tween-20 (TBS-T) and incubated overnight with 1:1000 dilutions of primary antibodies in 5% BSA in TBS-T. The membranes were washed three times with TBS-T, incubated for 1 h at room temperature with 1:10,000 dilutions of HRP-conjugated secondary antibodies (Cytiva Life Sciences) in 5% BSA in TBS-T, washed three times with TBS-T, washed once with TBS. Proteins were detected using ECL2 or PicoPLUS reagent (Thermo Fisher Scientific).

### Inside-out plasma membrane vesicle immunopurification and mass spectrometry

HCT116 Lck-mEGFP-3xHA cells were grown to ∼90% confluence in six 150 mm dishes. Lck-mEGFP-3xHA expression was induced by replacing the conditioned medium with DMEM supplemented with 10% FBS and 0.125 µg/ml doxycycline 24 h prior to the experiment. All subsequent manipulations were performed at 4°C. Each 150 mm dish was washed once with 5 ml of cold PBS supplemented with 1 mM EGTA then harvested by scraping into 5 ml of cold PBS containing 1 mM EGTA. The cells were collected by centrifugation at 300xg for 5 min, and the supernatant was discarded. The cell pellets were resuspended in 2 vol of organelle lysis buffer (1X TBS, 1 mM EGTA, 5 mM TCEP, 25 µM Calpain-I inhibitor and a protease inhibitor cocktail (see immunoblotting section)). The cell suspension was mechanically lysed by passing through a 25-gauge needle until ∼85% of cells were lysed as assessed by trypan blue exclusion. The lysed cells were centrifuged at 1,000xg for 15 min to sediment intact cells and nuclei, and the PNS was transferred to a 15 ml conical tube.

Beads from 200 µl of a magnetic Protein G Dynabead (Thermo Fisher Scientific) slurry were sedimented with a magnetic tube rack and resuspended in organelle lysis buffer. The bead slurry was divided evenly into four tubes and re-sedimented. The PNS was equally divided and divided between the four tubes. Tubes 1 and 2 received 5 µg of mouse IgG isotype antibody (Cell Signaling Technology G3A1), and tubes 3 and 4 received 5 µg of mouse anti-HA antibody (Proteintech 66006-2-Ig). Tubes 2 and 4 also received CaCl_2_ (2 mM final). The binding reactions were incubated on a rotating mixer for 15 min and washed three times with organelle lysis buffer. The wash buffer for tubes 2 and 4 was supplemented with 2 mM CaCl_2_. Each sample was eluted by heating at 55°C for 5 min in 65 µl of elution buffer (50 mM TEAB and 0.1% Rapigest (Waters)). An aliquot of each eluate (15 µl) was reserved for Sypro Ruby staining and immunoblot analysis. The remaining eluates from tubes 3 and 4 (three biological replicates each) were sent to the UC Berkeley Vincent J. Coates proteomics facility for label-free quantitative mass spectrometry. The mass spectrometry data were analyzed using PEAKS Studio.

### Genome-wide CRISPRi screens for plasma membrane repair

MDA-MB-231 cells expressing Zim3-KRAB-dCas9-P2A-Hygro (Addgene #188768) were transduced with the genome-wide CRISPRi library at a multiplicity of infection (MOI) of 0.2. After puromycin selection, nine 150 mm dishes of CRISPRi library cells at ∼80% confluence were washed once with 10 ml PBS and dissociated using 5 ml of accutase per dish. The cell suspension was pooled in a 50 ml conical tube and centrifuged at 300xg for 4 min. The cells were resuspended in 12 ml of TBS, divided equally into 3 tubes, and adjusted to 2 million cells/ml in each tube. Each tube received CaCl_2_, EGTA and/or PFO to achieve the following final concentrations: 2 mM CaCl_2_ (untreated), 0.78 ng/ml PFO and 2 mM EGTA (PFO + EGTA), or 3.125 ng/ml PFO and 2 mM CaCl_2_ (PFO + Ca^2+^). Cells were mixed by gentle inversion and incubated at 37°C for 20 min. Cells were sedimented at 300xg for 4 min and resuspended in 12 ml of conditioned medium. The cell suspensions were distributed equally into six 150 mm dishes per condition. After 36 h, the cells (∼25 million cells per condition) were harvested with accutase and stored at-80°C.

### Genomic DNA isolation

Cells from each CRISPRi screen condition were lysed in 5 ml of genomic DNA lysis buffer (20 mM Tris-HCl pH 8.0, 10 mM EDTA, 100 mM NaCl and 0.5% SDS) in a 15 ml conical tube. DNA was extracted by sequential phenol:chloroform:isoamyl alcohol (25:24:1, pH 8.0) and chloroform extractions, each performed by mixing until an emulsion formed, incubating for 5 min, and centrifuging at 5,000xg for 1 h. Between extractions, 2.5 ml of TE buffer was added to each organic phase for re-extraction, and the aqueous phases were pooled. DNA was precipitated from each combined aqueous phase (∼10 ml) by addition of sodium acetate (1 ml of 3 M, pH 5.2) and ethanol (20 ml) and stored overnight at 4°C.

Each sample was recovered by centrifugation at 5,000xg at 4°C for 1 h. The pellets were resuspended in the remaining supernatant (∼1 ml), transferred to a 1.5 ml tube, and centrifuged at 20,000xg at 4°C for 5 min. The pellets were washed twice with 1 ml of 70% ethanol by centrifugation at 15,000xg at room temperature for 5 min and allowed to air dry for 10 min.

The dried pellets were resuspended in 0.75 ml of 1X RNase ONE reaction buffer (Promega), and RNase ONE (3.75 µl; Promega) and RNase A (Thermo Fisher Scientific) were added and incubated at 37°C for 6 h. DNA was re-precipitated by the addition of sodium acetate (75 µl of 3 M, pH 5.2) and ethanol (1.5 ml), stored overnight at 4°C, and each sample was recovered as described above before final resuspension in 0.75 ml of 2.5 mM Tris-HCl pH 8.0.

### CRISPRi library preparation and sequencing

A PCR reaction was assembled with the following components: 20 µl of oJR441 reverse primer (100 µM stock), 20 µl of indexed forward primer (100 µM stock), 1 ml of NEBNext Ultra II Q5 MasterMix, and 200 µg of genomic DNA. Each reaction was adjusted to 2 ml with nuclease-free water and distributed into 100 µl aliquots. PCR amplification was performed as follows: (1) 98°C for 30 sec; (2) 98°C for 10 sec; (3) 63°C for 75 sec; (4) steps 2-3 repeated 21 times; (5) 72°C for 5 min. PCR amplicons were purified using a 0.5-0.65X double SPRI size selection (Beckman Coulter). The libraries were spiked with 5% PhiX and sequenced on a NextSeq P2 100PE flow cell (400 million paired-end reads, 25 cycles) using custom sequencing primers.

- oJR441 reverse primer sequence: CAAGCAGAAGACGGCATACGAGATGCGGCCGGCTGTTTCCAGCTTAGCTCTTAAA
- Indexed forward primer sequence: AATGATACGGCGACCACCGAGATCTACACnnnnnnnnCGCGTATCCCTTGGAGAACCACC
- Read 1 primer sequence: CGCGTATCCCTTGGAGAACCACCTTGTTGG
- Read 2 primer sequence: GCGGCCGGCTGTTTCCAGCTTAGCTCTTAAAC
- Index: CCAACAAGGTGGTTCTCCAAGGGATACGCG

### CRISPRi screen analysis

A custom MATLAB (MathWorks) script was used to count the number of reads aligning to both sgRNAs for each gene. Genes with read counts in the bottom 20^th^ percentile were excluded from analysis to remove genes whose depletion affected cell growth and viability.

### Live-cell microscopy and laser ablation

Images were acquired using an LSM900 confocal microscope (ZEISS) using confocal mode, a 63× Plan-Apochromat, NA 1.40 objective, and a heated, CO_2_-controlled chamber. Cells were seeded in 35 mm collagen-coated glass-bottom dishes (MatTek) and imaged in Fluorobrite DMEM supplemented with 10% FBS and 2.5 µM FM4-64 (Biotium). For laser ablation experiments, a 1 µm x 1 µm square region of interest was positioned over the edge of a cell not adjacent to another cell and ablated for 100 iterations using all lasers at 100% power. Protein recruitment was quantified in Zen 3.1 (Zeiss) by measuring fluorescence intensities within a box that encapsulated the repair cap. Fluorescence intensities of an adjacent, non-ablated area of the membrane was measured and used for a frame-by-frame background subtraction. Recruitment was plotted as the background-corrected change in fluorescence intensity divided by the fluorescence intensity at the damage site prior to ablation.

For the laser ablation experiments in Figure 3C, the cells were washed once with TBS and imaged in TBS supplemented with 2.5 µM FM4-64 and either 2 mM CaCl_2_ or 2 mM EGTA.

### Immunofluorescence microscopy

U2-OS cells grown to ∼70% confluence on 12 mm glass coverslips (Corning) were treated with vehicle or 25 ng/ml PFO for 5 min at 37°C. The cells were washed once with PBS, fixed in 4% EM-grade paraformaldehyde (Electron Microscopy Science), washed three times with PBS, and permeabilized and blocked in IF blocking buffer (2% FBS and 0.02% saponin in PBS) for 30 min at room temperature. Coverslips were transferred to a humidity chamber, incubated with primary antibodies (1:100 dilution in IF blocking buffer) for 1 h at room temperature, washed three times with PBS, incubated with fluorophore-conjugated secondary antibodies (1:500 dilution in IF blocking buffer) for 1 h at room temperature, and washed three times with PBS. The coverslips were then mounted overnight in ProLong Gold antifade mountant with DAPI (Thermo Fisher Scientific) and sealed with clear nail polish. Images were acquired using an LSM900 confocal microscope using Airyscan 2 mode and a 63× Plan-Apochromat, NA 1.40 objective.

### RNA interference

U2-OS cells were suspended using accutase and transfected using Lipofectamine RNAiMAX (Thermo Fisher Scientific) according to the manufacturer’s instructions. The cell suspension was diluted in conditioned medium to a final concentration of 1.5 × 10^5^ cells/ml, 20 nM total siRNA and 0.3% (v/v) transfection reagent, dispensed into individual wells of a poly-D-lysine-coated six-well plate and incubated overnight at 37°C. Conditioned medium was replaced after 18-24 h. At 48 h post-transfection, cells were suspended using accutase, reseeded at an appropriate density for laser ablation, cell viability, or SYTOX Green influx experiments and incubated at 37°C for an additional 24 h. The following siRNAs were purchased from Qiagen: ANXA11 (Hs_ANXA11_6), SRI (Hs_SRI_10) and a non-targeting control (1027310).

### Cell viability assays

U2-OS cells were transfected with siRNAs as described above, seeded in a poly-D-lysine-coated 24-well plate and grown to 70% confluence. At 72 h post-transfection, cells were treated with vehicle or 25 ng/ml PFO for 30 min in 200 μl of Fluorobrite DMEM. Fluorobrite DMEM was replaced with 200 µl of fresh DMEM supplemented with 10% FBS, and the cells were allowed to recover for 1 h at 37°C. After the recovery period, 200 µl of Fluorobrite DMEM supplemented with 0.5 mg/ml MTT reagent was added to each well and incubated at 37°C for 1 h. The MTT-containing medium was removed, replaced with 200 µl of DMSO and incubated for 5 min. An aliquot of each DMSO suspension (100 µl) was transferred to a 96-well plate, and the absorbance was measured at 570 nm. Cell viability was calculated by subtracting the absorbance of a DMSO blank from each sample measurement.

### Measurement of plasma membrane permeabilization by PFO

U2-OS cells were transfected with siRNAs as described above, seeded in 35 mm collagen-coated glass-bottom dishes and grown to 70% confluence. At 72 h post-transfection, cells were washed with PBS, placed in a 37°C, 5% CO_2_ chamber on an LSM900 confocal microscope system and incubated with Fluorobrite DMEM supplemented with 10% FBS, 25 ng/ml PFO and 2.5 μM SYTOX Green. SYTOX Green uptake was measured using the 20× Plan-Apochromat, NA 0.8 objective with images acquired every 10 sec for 10 min.

### Recombinant protein purification

Recombinant proteins were purified as described in (59) with the following modifications. Recombinant PFO was expressed in *E. coli* Rosetta2(DE3)pLysS cells. A pre-culture (5 ml) was grown overnight at 37°C and diluted into a 500 ml culture. The culture was incubated at 37°C until the OD_600_ reached ∼0.6, and protein expression was induced by addition of 50 µM IPTG overnight at 30°C. The cells were harvested by centrifugation at 5,000xg for 10 min in a swing-bucket H-6000A rotor, and the cell pellet was stored at-80°C until use. All subsequent manipulations were performed at 4°C. The cell pellet was resuspended in 5 ml of Ni-NTA lysis buffer 1 (50 mM HEPES pH 7.4, 150 mM NaCl, 10 mM imidazole pH 7.4 and a protease inhibitor cocktail) and sonicated 8 times (5 sec on, 10 sec off, 15% amplitude). The lysate was clarified by centrifugation at 5,000xg for 10 min in a fixed angle FIBERlite F21-8×50y rotor, and the supernatant was added to 1 ml of pre-equilibrated HisPur^TM^ Ni-NTA Resin and incubated on a rotating mixer for 1 h. The beads were washed three times with Ni-NTA wash buffer 1 (50 mM HEPES pH 7.4, 250 mM NaCl, 10 mM imidazole pH 7.4) and eluted with Ni-NTA elution buffer 1 (50 mM HEPES pH 7.4, 150 mM NaCl, 300 mM imidazole pH 7.4). The eluate was diluted with dilution buffer (50 mM HEPES pH 7.4, 150 mM NaCl, 10 mM DTT) to a final protein concentration of 1 mg/ml, snap-frozen in liquid nitrogen and stored at-80°C until use.

Recombinant sorcin and ANXA2 were expressed in *E. coli* Rosetta2(DE3)pLysS cells. Pre-cultures (30 ml) were grown overnight at 37°C and diluted into 3000 ml cultures. The cultures were incubated at 37°C until the OD_600_ reached ∼0.6, and protein expression was induced by addition of 0.2 mM IPTG for 4 h at 37°C. The cells were harvested by centrifugation at 5,000xg for 10 min in a swing-bucket H-6000A rotor, and cell pellets were stored at-80°C until use. All subsequent manipulations were performed at 4°C. The cell pellets were resuspended in 30 ml of Ni-NTA lysis buffer 2 (20 mM Tris-HCl pH 8.0, 300 mM NaCl, 10 mM imidazole pH 8.0 and a protease inhibitor cocktail) and sonicated 5 times (5 sec on, 15 sec off, 20% amplitude). Each lysate was clarified by centrifugation at 20,000xg for 15 min in a fixed angle FIBERlite F21-8×50y rotor, and the supernatant fractions were applied to gravity flow columns containing 1 ml of pre-equilibrated HisPur^TM^ Ni-NTA Resin (Thermo Fisher Scientific). Each column was washed with three column volumes of Ni-NTA wash buffer 2 (20 mM Tris-HCl pH 8.0, 300 mM NaCl, 25 mM imidazole pH 8.0), eluted with Ni-NTA elution buffer 2 (20 mM Tris-HCl pH 8.0, 300 mM NaCl, 300 mM imidazole pH 8.0). Eluted protein at 0.5 mg/ml was aliquoted, snap-frozen in liquid nitrogen and stored at-80°C until use. Recombinant His_6_-ANXA11 was purchased from Bio-Techne.

### Endogenous sorcin immunoprecipitation

HCT116 wild-type cells were grown to ∼90% confluence in four 150 mm dishes. All subsequent manipulations were performed at 4°C. Each dish was washed once with 5 ml of cold PBS supplemented with 1 mM EGTA and harvested by scraping into 5 ml of cold PBS containing 1 mM EGTA. Cells were pooled, collected by centrifugation at 300xg for 5 min, and the supernatant was discarded. The cell pellet was resuspended in 2.5 ml of IP buffer (1X TBS, 1 mM EGTA, 0.5% TX-100 and a protease inhibitor cocktail) and incubated for 10 min. The lysed cells were centrifuged at 15,000xg for 10 min to sediment insoluble material, and the supernatant was transferred to a new tube. An aliquot (50 μl) of the supernatant was retained for an input measurement, and the remaining lysate was distributed equally to four tubes.

Tubes 1 and 2 received 5 µg of mouse IgG isotype antibody (Cell Signaling Technology G3A1), and tubes 3 and 4 received 5 µg of mouse anti-sorcin antibody (Santa Cruz; sc-100859). Tubes 2 and 4 also received CaCl_2_ (2 mM final). Pre-equilibrated Protein A/G agarose beads (50 μl) (Santa Cruz Biotechnology) was added to each tube, and the binding reactions were incubated on a rotating mixer overnight. The beads were then washed three times with IP buffer. The wash buffer for tubes 2 and 4 was supplemented with 2 mM CaCl_2_. Each sample was eluted in 1X Laemmli buffer (non-reducing) for immunoblot analysis.

### FLAG immunoprecipitation

HCT116 wild-type cells were grown to ∼90% confluence in ten 150 mm dishes. All subsequent manipulations were performed at 4°C. Each dish was washed once with 5 ml of cold PBS supplemented with 1 mM EGTA and harvested by scraping into 5 ml of cold PBS containing 1 mM EGTA. Cells were pooled, collected by centrifugation at 300xg for 5 min, and the supernatant was discarded. The cell pellet was resuspended in 2 vol of organelle lysis buffer, and the cell suspension was mechanically lysed by passing through a 25-gauge needle until ∼85% of cells were lysed as assessed by trypan blue exclusion. The lysed cells were centrifuged at 1,000xg for 15 min to sediment intact cells and nuclei, and the PNS was centrifuged at 100,000xg for 30 min in an SW55 rotor. The supernatant (cytosol fraction) was collected carefully without disturbing the membrane pellet and diluted 1:2 in organelle lysis buffer.

In parallel, 150 µl of anti-FLAG agarose beads (Chromotek ffa-20) were sedimented at 1,000xg for 2 min and resuspended in 600 µl of organelle lysis buffer. The resuspended beads were divided equally into six tubes. FLAG (3x) peptide, SRI^FL^-FLAG and SRI^Δ1-32^-FLAG baits (4 µM each) were added to anti-FLAG beads in a 200 µl reaction. Bait proteins were bound to the anti-FLAG beads for 1 h at 4°C. The beads were washed three times with organelle lysis buffer and sedimented at 1,000xg for 2 min. The purified cytosol was equally divided and dispensed between the six tubes. CaCl_2_ (2 mM final) was added to the indicated tubes. The binding reactions were incubated on a rotating mixer for 2 h. The beads were washed three times with organelle lysis buffer. The wash buffer for the Ca^2+^ immunoprecipitations was supplemented with 2 mM CaCl_2_. Each sample was eluted in 1X Laemmli buffer (non-reducing) for immunoblot analysis.

### Liposome production

Liposomes were generated by mixing lipids in a glass vial at the following molar ratios: 38:20:20:3:0.5:18.5 DOPC:POPE:DOPS:PI(4,5)P_2_:TexasRed-PE:cholesterol (1 µmol total lipid) in 500 µl of 5% methanol and 95% chloroform. The lipid mixture was placed on a heat block preheated to 55°C, and the organic solvent was evaporated under vacuum for 30 min. Dried lipids were resuspended in 500 µl of degassed TBS and incubated at 55°C with intermittent vortexing for 15 min. The lipid solution was extruded 11 times through a 200-nm filter.

### Buoyant density gradient flotation

Buoyant density gradient flotation assays were performed as described in (60) with the following modifications. Binding reactions (60 µl) consisted of liposomes (20 µl), purified proteins and CaCl_2_ (2 mM final) in TBS. The reactions were incubated at room temperature for 15 min, mixed with 60 µl of 60% sucrose in TBS supplemented with 2 mM CaCl_2_ (for a final concentration of 30% sucrose), and sequentially overlaid with 100 µl of 25% sucrose in TBS supplemented with 2 mM CaCl_2_ and 20 µl of TBS supplemented with 2 mM CaCl_2_. The sucrose step gradients were centrifuged in a TLA100 rotor (Beckman) at 100,000 rpm (436,000xg) for 10 min at 4°C. Buoyant liposomes were collected from the top of the gradient, and the bound proteins were analyzed by SDS-PAGE and immunoblotting.

## Statistical analysis

Statistical analyses were performed using GraphPad Prism 10. Unpaired two-tailed Student’s *t*-tests were used for comparisons between two groups, and one-or two-way analysis of variance (ANOVA) tests were used for comparisons among multiple groups. The statistical test used for each experiment is detailed in the figure legends.

## Acknowledgements

We dedicate this work to Bob Lesch, our lab manager for the past several decades who was tragically taken from us by an accident in 2021. We thank all members of the Schekman lab for thoughtful discussions during the preparation of this manuscript. We thank Claire Goul and Roberto Zoncu for providing plasmids to generate the ALIX-mNG and CHMP4B-mNG cell lines. We thank Sankalp Shukla and Jim Hurley for providing the purified ALIX protein. We also thank our current lab manager, Nam Che, Rob Maxwell from the UC Berkeley Vincent J. Coates Proteomics Facility and Alison Killilea from the UC Berkeley Cell Culture Facility. This work used the Vincent J. Proteomics/Mass Spectrometry Laboratory at UC Berkeley, supported in part by NIH S10 Instrumentation Grant S10RR025622.

## Author Contributions

J.M. Ngo, J.K. Williams and R. Schekman conceptualized the study. J.M. Ngo, J.K. Williams and A. Murugupandiyan carried out the research experiments. J.M. Ngo, J.K. Williams and R. Schekman analyzed the data, prepared the original draft and revised the manuscript.

## Movies

**Movie 1. Laser ablation of U2-OS SRI-mNG cells.** U2-OS cells stably expressing SRI-mNG were subjected to laser ablation. Green: SRI-mNG; Magenta: FM4-64. Image times are relative to the first image taken after laser ablation. Total imaging time: 5 min. Time between acquisitions: 3 s. Scale bars: 5 μm. See corresponding Figure 3A.

**Movie 2. Laser ablation of U2-OS SRI-mNG cells in the presence of extracellular Ca^2+^.** U2-OS cells stably expressing SRI-mNG were subjected to laser ablation in the presence of extracellular Ca^2+^. Green: SRI-mNG; Magenta: FM4-64. Image times are relative to the first image taken after laser ablation. Total imaging time: 5 min. Time between acquisitions: 3 s. Scale bars: 5 μm. See corresponding Figure 3C.

**Movie 3. Laser ablation of U2-OS SRI-mNG cells in the absence of extracellular Ca^2+^.** U2-OS cells stably expressing SRI-mNG were subjected to laser ablation in the absence of extracellular Ca^2+^. Green: SRI-mNG; Magenta: FM4-64. Image times are relative to the first image taken after laser ablation. Total imaging time: 5 min. Time between acquisitions: 3 s. Scale bars: 5 μm. See corresponding Figure 3C.

**Movie 4. Laser ablation of U2-OS SRI-mNG siNT cells.** U2-OS cells stably expressing SRI-mNG were transfected with a non-targeting siRNA (siNT) and subjected to laser ablation. Green: SRI-mNG; Magenta: FM4-64. Image times are relative to the first image taken after laser ablation. Total imaging time: 5 min. Time between acquisitions: 3 s. Scale bars: 5 μm. See corresponding Figure 5D.

**Movie 5. Laser ablation of U2-OS SRI-mNG siANXA11 cells.** U2-OS cells stably expressing SRI-mNG were transfected with an ANXA11-targeting siRNA (siANXA11) and subjected to laser ablation. Green: SRI-mNG; Magenta: FM4-64. Image times are relative to the first image taken after laser ablation. Total imaging time: 5 min. Time between acquisitions: 3 s. Scale bars: 5 μm. See corresponding Figure 5D.

**Movie 6. Laser ablation of U2-OS ANXA11-mSc siNT cells.** U2-OS cells stably expressing ANXA11-mSc were transfected with a non-targeting siRNA (siNT) and subjected to laser ablation. Green: ANXA11-mSc; Magenta: FM4-64. Image times are relative to the first image taken after laser ablation. Total imaging time: 5 min. Time between acquisitions: 3 s. Scale bars: 5 μm. See corresponding Figure 5F.

**Movie 7. Laser ablation of U2-OS ANXA11-mSc siSRI cells.** U2-OS cells stably expressing ANXA11-mSc were transfected with a sorcin-targeting siRNA (siSRI) and subjected to laser ablation. Green: ANXA11-mSc; Magenta: FM4-64. Image times are relative to the first image taken after laser ablation. Total imaging time: 5 min. Time between acquisitions: 3 s. Scale bars: 5 μm. See corresponding Figure 5F.

**Movie 8. Laser ablation of U2-OS ALIX-mNG siNT cells.** U2-OS cells stably expressing ALIX-mNG were transfected with a non-targeting siRNA (siNT) and subjected to laser ablation. Green: ALIX-mNG; Magenta: FM4-64. Image times are relative to the first image taken after laser ablation. Total imaging time: 5 min. Time between acquisitions: 3 s. Scale bars: 5 μm. See corresponding Figure 6A.

**Movie 9. Laser ablation of U2-OS ALIX-mNG siSRI cells.** U2-OS cells stably expressing ALIX-mNG were transfected with a sorcin-targeting siRNA (siSRI) and subjected to laser ablation. Green: ALIX-mNG; Magenta: FM4-64. Image times are relative to the first image taken after laser ablation. Total imaging time: 5 min. Time between acquisitions: 3 s. Scale bars: 5 μm. See corresponding Figure 6A.

**Movie 10. Laser ablation of U2-OS ALIX-mNG siANXA11 cells.** U2-OS cells stably expressing ALIX-mNG were transfected with an ANXA11-targeting siRNA (siANXA11) and subjected to laser ablation. Green: ALIX-mNG; Magenta: FM4-64. Image times are relative to the first image taken after laser ablation. Total imaging time: 5 min. Time between acquisitions: 3 s. Scale bars: 5 μm. See corresponding Figure 6A.

**Movie 11. Laser ablation of U2-OS CHMP4B-mNG siNT cells.** U2-OS cells stably expressing CHMP4B-mNG were transfected with a non-targeting siRNA (siNT) and subjected to laser ablation. Green: CHMP4B-mNG; Magenta: FM4-64. Image times are relative to the first image taken after laser ablation. Total imaging time: 5 min. Time between acquisitions: 3 s. Scale bars: 5 μm. See corresponding Figure 6C.

**Movie 12. Laser ablation of U2-OS CHMP4B-mNG siSRI cells.** U2-OS cells stably expressing CHMP4B-mNG were transfected with a sorcin-targeting siRNA (siSRI) and subjected to laser ablation. Green: CHMP4B-mNG; Magenta: FM4-64. Image times are relative to the first image taken after laser ablation. Total imaging time: 5 min. Time between acquisitions: 3 s. Scale bars: 5 μm. See corresponding Figure 6C.

**Movie 13. Laser ablation of U2-OS CHMP4B-mNG siANXA11 cells.** U2-OS cells stably expressing CHMP4B-mNG were transfected with an ANXA11-targeting siRNA (siANXA11) and subjected to laser ablation. Green: CHMP4B-mNG; Magenta: FM4-64. Image times are relative to the first image taken after laser ablation. Total imaging time: 5 min. Time between acquisitions: 3 s. Scale bars: 5 μm. See corresponding Figure 6C.

## Tables

Table 1. Quantitative proteomics of inside-out PM vesicles in the absence or presence of Ca^2+^.

Table 2. Genome-wide CRISPR interference screens for cell survival after PM damage.

